# The impact of downsampling on data quality, univariate measurement and multivariate pattern analysis in event-related potential research

**DOI:** 10.64898/2025.12.24.696322

**Authors:** Guanghui Zhang, Xinran Wang, Ying Xin, Fengyu Cong, Weiqi He, Wenbo Luo

## Abstract

The choice of sampling rate is a critical preprocessing step in event-related potential (ERP) research, yet its impact on different analytic approaches remains underexplored. In this study, we systematically evaluated how downsampling affects data quality measured via Standardized Measurement Error (SME), conventional univariate ERP metrics (mean amplitude, peak amplitude, peak latency, and 50% area latency), and multivariate pattern analysis (MVPA; decoding). We analyzed seven commonly studied ERP components: P3, N400, N170, N2pc, mismatch negativity, error-related negativity, and lateralized readiness potential collected from neurotypical young adults. Results showed that both amplitude- and latency-based scores were significantly affected by sampling rate changes, particularly for 50% area latency that presented increased SME (reduced data quality) at lower sampling rates. Amplitude-based measures and their corresponding effect sizes were more influenced by sampling rate variations, whereas latency-based measures were comparatively stable across different temporal resolutions. In contrast, multivariate decoding performance remained highly robust, with decoding accuracy and effect sizes showing minimal variation even at the lowest sampling rate (e.g., 64 Hz). Overall, these findings suggest that caution should be exercised when employing lower sampling rates for data quality and conventional univariate ERP analyses. Nevertheless, lower sampling rates (e.g., 64 Hz) may still be appropriate for decoding analyses, particularly in studies where precise temporal resolution is not critical. For researchers analyzing ERP data with similar components, noise levels, and participant populations as in this study, following these recommendations should yield robust statistical power.

## 1. Introduction

The selection of an appropriate sampling rate is an important preprocessing step in electroencephalographic (EEG) research (Pedroni et al., 2019). In event-related potential (ERP) studies, where millisecond-level temporal precision is often required, the sampling rate determines how finely the neural time course is captured. While traditional recommendations suggest sampling rates of 200–1200 Hz are sufficient for most ERP components (Luck, 2014), these guidelines are typically based on signal-processing heuristics (e.g., the Nyquist theorem), historical precedent, or lab lore, rather than systematic empirical evidence comparing different analytic approaches. As a result, researchers may underestimate how sampling rate choices affect not only data quality of the signals, but also outcomes of both univariate and multivariate pattern analyses (MVPA; decoding).

In practical applications, downsampling is often employed before computing classic ERP metrics, such as mean amplitude, peak amplitude, 50% area latency, and peak latency, to reduce data size, computational load, and increase statistical power. Clayson and colleagues (2013) investigated the impact of downsampling from 2000 Hz to 1000, 500, and 250 Hz on amplitude and latency estimates using both real and simulated ERP data. They found that mean amplitude and area latency remained relatively robust and statistically efficient across different sampling rates, whereas peak amplitude and peak latency were more susceptible to noise and estimation error. Similarly, Jing and Takigawa (2000) reported that low sampling rates can distort nonlinear signal characteristics, such as correlation dimension, highlighting the potential for lower temporal resolution to introduce subtle but meaningful signal degradation. Despite these findings, sampling rate settings vary widely across published studies, even within the same research domain. For example, Beres (2017) ERP-based language studies employed sampling rates ranging from 250 Hz to 512 Hz. Moreover, formal recommendations also differ: Luck (2014) suggested that sampling rates between 200 and 1200 Hz are generally sufficient for most cognitive and affective neuroscience experiments, but such recommendations may not adequately account for modern analytic techniques (e.g., decoding).

Beyond traditional ERP metrics, decoding analysis has gained prominence as a powerful approach for extracting information from ERP signals (Ashton et al., 2022; Grootswagers et al., 2017). Decoding analysis leverages spatiotemporal patterns in the EEG to classify stimuli, predict behavioral responses, or decode latent cognitive states. Often regarded as more sensitive than univariate methods (King & Dehaene, 2014), decoding analysis also presents new challenges related to temporal resolution and data dimensionality. Several studies have examined how downsampling or temporal smoothing affects decoding performance. While some report minimal effects of moderate downsampling (e.g., Davis et al., 2018; Meribout et al., 2023), others suggest that lower sampling rates, even as low as 50 Hz, can enhance classification accuracy (Brandmeyer et al., 2013; Grootswagers et al., 2017). Despite these seemingly contradictory findings, there is still no clear consensus on whether and how sampling rate influences decoding performance. Consequently, sampling rates in MVPA studies vary widely from 50 Hz to 2048 Hz depending on the experimental design, data quality, and population characteristics (Brandmeyer et al., 2013; Leong et al., 2025; Liu et al., 2025; Marsicano et al., 2024; Sarrett & Toscano, 2024; Zhang & Luck, 2025).

In addition to influencing signal interpretability and analysis outcomes, sampling rate also affects measurement precision. One sensitive index of data quality is the Standardized Measurement Error (SME; Luck et al., 2021), which quantifies trial-to-trial variability relative to the signal of interest. While SME has been widely applied to assess the effects of filter settings and different scoring methods (Zhang, Garrett, & Luck, 2024b; Zhang & Luck, 2023), few studies have directly examined how it varies as a function of sampling rate. Notably, Jing & Takigawa (2000) demonstrated that low sampling rates can inflate nonlinear estimates such as correlation dimension, suggesting that lower temporal resolution introduces subtle signal distortions that may affect precision.

Although some recent papers have provided valuable commentary on the interplay between downsampling and statistical sensitivity (e.g., Brandmeyer et al., 2013; Clayson et al., 2013) through a single study, no comprehensive study to date has systematically examined how sampling rate impacts data quality, ERP measurements, and decoding performance within a large set of datasets.

### 1.1 The aims of present study

In this study, we systematically evaluated the impact of sampling rate on three critical aspects of ERP analysis: (1) signal quality, assessed by SME, (2) traditional univariate metrics including mean amplitude, peak amplitude, peak latency, and 50% area latency, and (3) multivariate decoding accuracy using time-resolved MVPA. To ensure broad applicability, we conducted these analyses across seven widely studied ERP components [P3, N400, N170, N2pc, mismatch negativity (MMN), error-related negativity (ERN), and lateralized readiness potential (LRP)] using a large dataset collected from neurotypical young adults. This component-wise, multidimensional approach allows for a generalizable assessment of how temporal resolution interacts with both cognitive processes and analysis strategies.

By comparing results across a range of downsampled datasets (64, 128, 256, 512, and 1024 Hz), we provided empirical guidance for researchers aiming to balance data efficiency with analytic rigor. Our findings offer nuanced insights into when lower sampling rates may suffice, particularly for amplitude-based or classification-based analyses, and when higher sampling resolution is necessary to preserve temporal fidelity, especially for latency-based metrics.

## 2 Methods

In this study, we conducted all analyses using publicly available EEG datasets which have been preprocessed to the epoch data from ERP CORE (Compendium of Open Resources and Experiments; Kappenman et al., 2021). In the following sections, we will briefly describe the experiment process, data preprocessing pipelines, outline the data quality measurement, univariate analysis, and decoding analysis separately.

The data analysis procedures were implemented with MATLAB 2023a (MathWorks Inc), using EEGLAB v2024.0 (Delorme & Makeig, 2004) combined with ERPLAB v12.00 (Lopez-Calderon & Luck, 2014).

The raw datasets were available at: https://doi.org/10.18115/D5JW4R. The datasets preprocessed with optimized ICA procedure and Matlab scripts used in this study were available at https://osf.io/v5suh/.

### 2.1 Participants

All datasets included 40 college students with self-reported normal visual acuity and no history of major neurological disorders (N = 40) from University of California, Davis community.

### 2.2 Experimental paradigms

The ERP CORE datasets were obtained from six different experiment paradigms, which elicited seven different ERP components. Participants completed all six tasks in one session, with each task presumably lasting ten minutes.

As shown in Fig.1a, we used a face perception paradigm to elicit the N170 component. The experiment included four types of stimuli: face, car, scrambled face, and scrambled car. On each trial, a type of stimuli was randomly presented in the center of the screen. Participants were instructed to respond to the stimulus as quickly as possible, indicating whether the stimulus was a “texture” (scrambled face or scrambled car) or an “object” (face or car).

**Figure 1.**
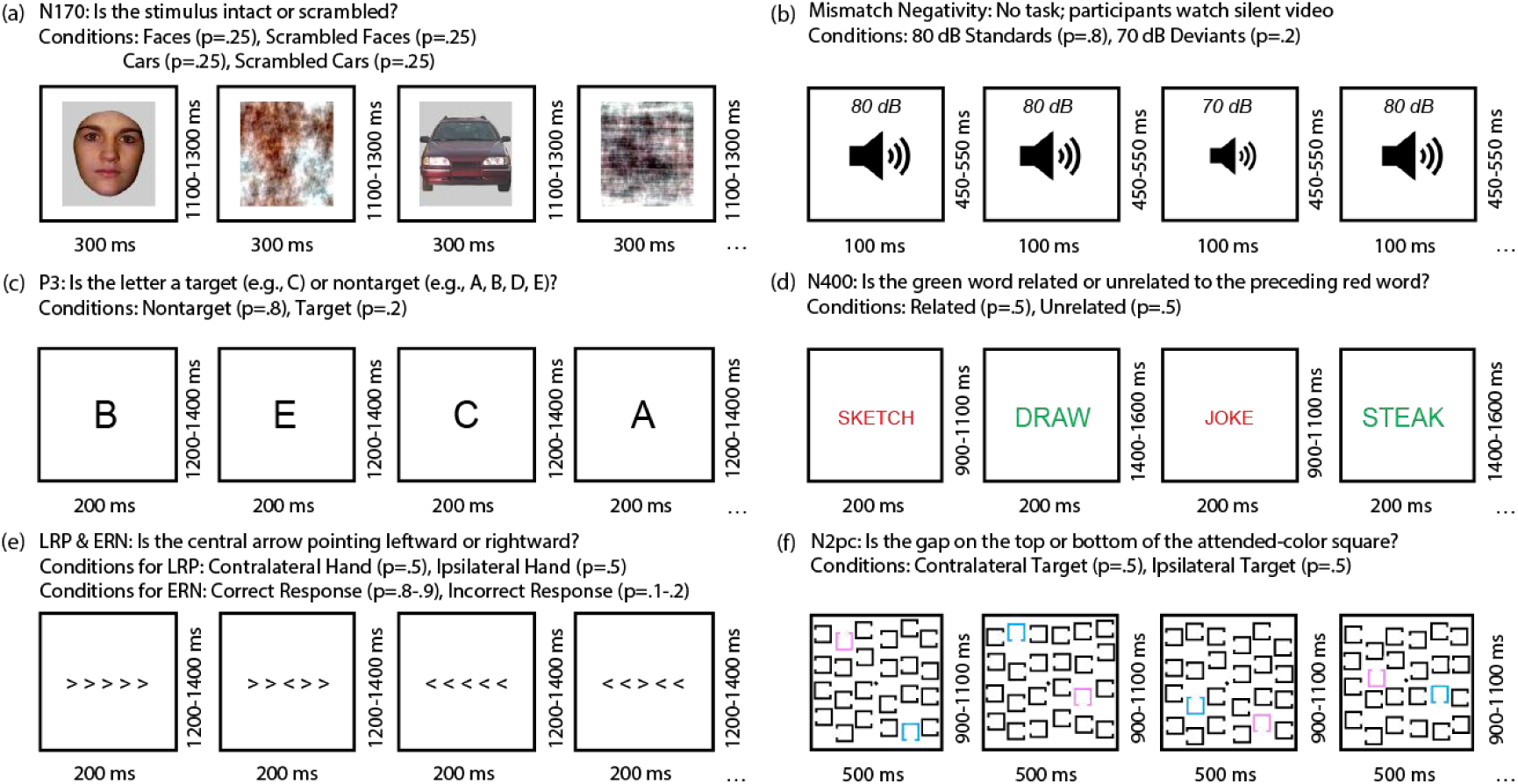
Examples of multiple trials from each of the six experimental paradigms are shown: (a) Face perception paradigm used to evoke the N170 component. Stimuli were randomly drawn from four equally probable categories: face, scrambled face, car, and scrambled car. Participants judged whether each image depicted an intact object (face or car) or a texture (scrambled versions). The current analysis focuses exclusively on face and car trials. (b) Passive auditory oddball paradigm designed to elicit mismatch negativity (MMN). On each trial, participants heard either a standard tone (80 dB; p = .8) or a deviant tone (70 dB; p = .2). These auditory stimuli were task-irrelevant, and participants viewed a silent video throughout the task. (c) Active visual oddball paradigm designed to elicit the P3 component. Letters A, B, C, D, and E were presented in random order (each with a probability of .2). In each block, one letter served as the target (20% of trials), while the remaining four letters as nontargets (80%). Participants were instructed to classify each stimulus as either a target or a nontarget. (d) Word pair judgment paradigm used to elicit the N400 component. A red prime word was followed by a green target word. Participants indicated whether the target was semantically related (50% of trials) or unrelated (50%) to the prime. (e) Flanker task used to elicit both the lateralized readiness potential (LRP) and the error-related negativity (ERN). Participants responded to the direction of the central arrow while ignoring flanking arrows that could be congruent or incongruent. (f) Simple visual search task designed to elicit the N2pc component. One color (pink or blue) was assigned as the target at the beginning of each block. On each trial, participants reported whether the gap in the target-colored square was located at the top or bottom. The figure was adopted from Kappenman et al., 2021.

A passive auditory oddball paradigm was used to elicit the mismatch negativity (MMN; Fig. 1 b). Participants were presented with 80 dB standard stimuli (p = .8) and 20 dB deviation stimuli (p = .2). They were instructed to ignore auditory stimuli while watching a silent video.

An active visual oddball paradigm was used to elicit the P3 component (Fig.1 c). Participants were presented with one of five letters randomly (A, B, C, D, and E; p = .2 for each). Noted that one of the letters was designated as the target for one block and the others were nontargets. Participants were instructed to press one button if the presented stimulus was the target and another button if it was a non-target.

To elicit the N400 component (Fig.1 d), a word pair judgment paradigm was employed. Each trial consisted of a red prime word followed by a green target word. Participants were asked to judge whether the target word was semantically related or unrelated to the prime word and respond by pressing the corresponding button.

The Eriksen flanker’s paradigm was used to obtain the lateralized readiness potential (LRP) and the error-related negativity (ERN; Fig. 1e). In each trial, Participants identified the direction of a central arrowhead surrounded by congruent or incongruent arrowheads and they were asked to respond by pressing a button with their left or right hand accordingly.

To elicit the N2pc component (Fig.1 f), a visual search paradigm was used. Each stimuli array contained one pink square, one blue square, and 22 black squares. Either the pink or blue square served as the target stimulus. Participants were asked to identify the specific location (top or bottom) of the target color block gap and press the corresponding button. In all tasks, participants were instructed to maintain their gaze on the central fixation point throughout the trials.

### 2.3 EEG recording and preprocessing pipeline

All paradigms of EEG data were recorded using a Biosemi ActiveTwo recording system (Biosemi B.V.) with a sampling rate of 1024 Hz, which used a fifth-order sinc antialiasing filter with a half-power cutoff at 204.8 Hz. Recordings were obtained from 30 scalp sites (FP1/2, Fz/3/4/7/8, FCz/3/4, Cz/3/4/5/6, CPz, Pz/3/4/7/8, PO3/4/7/8/9/10, Oz/1/2) and electrooculogram (EOG) electrodes which are positioned beside the eyes and beneath the right eye.

For offline analyses, the data were first referenced to the average of P9 and P10 electrodes sites which were located near left and right mastoids. An exception was made for N170, for which the average reference across all electrodes was used. We then applied a noncausal Butterworth high-pass filter with 0.1 and low-pass filter with 20Hz to the data (12 dB/oct roll-off was used for both filters) (Luck, 2014; Zhang, Garrett, & Luck, 2024a, 2024b). We used 20Hz to avoid frequency leakage when low sampling rate was applied (e.g, 64Hz)^1^. We then performed independent component analysis (ICA) on the data to correct artifacts, such as blinks and eye movements with optimal pipeline as used in previous studies (Dimigen, 2020; Luck, 2022; Zhang, Garrett, Simmons, et al., 2024). To examine whether downsampling has a significant effect on data quality, univariate analysis and decoding analysis, we therefore resampled the data using six different sampling rates separately with EEGLAB function: 1024 Hz, 512 Hz, 256 Hz, 128 Hz, and 64 Hz. We then segmented and performed baseline correction on resampled data using a specific time window as shown in Tab. 1. The bad channels were then interpolated using EEGLAB’s spherical spline interpolation algorithm as the previous studies (Zhang, Carrasco, et al., 2024; Zhang, Garrett, Simmons, et al., 2024). Note that, for univariate analysis, rejection flags from the original ERP CORE datasets were applied to the present data, independent of the sampling rate settings for data quality and conventional univariate analysis. The marked epochs were then excluded from further analyses. However, we did not reject trials with artifacts for the decoding analysis, as previous research has shown that artifact rejection does not significantly affect decoding performance (Zhang & Luck, 2025).

**Table 1.** Epoch window, baseline period, and measurement window used for each experiment.

| Experiment | Epoch (ms) | Baseline Period (ms) | Measurement window (ms) | Electrode site |
| --- | --- | --- | --- | --- |
| N170 | -200 to 800 | -200 to 0 | 110 to 150 | PO8 |
| MMN | -200 to 800 | -200 to 0 | 125 to 225 | FCz |
| P3 | -200 to 800 | -200 to 0 | 300 to 600 | Pz |
| N400 | -200 to 800 | -200 to 0 | 300 to 500 | CPz |
| ERN | -600 to 400 | -400 to -200 | 0 to 100 | FCz |
| LRP | -800 to 200 | -800 to -600 | -100 to 0 | C3/C4 |
| N2pc | -200 to 800 | -200 to 0 | 200 to 275 | PO7/PO8 |

### 2.4 Univariate analysis

In this study, we computed mean amplitude, peak amplitude, 50% area latency and peak latency for different sampling rates separately. We used the time windows and component polarities specified by each experimental paradigm in the ERP CORE datasets (Tab. 1). This allowed us to calculate the amplitude and latency within a reasonable range.

For each sampling rate, we identified mean amplitude by averaging voltages for time points within the measurement window (specified in Tab. 1) for individual participants separately. Peak amplitude was identified as the maximum positive voltage for the P3 component, or the maximum negative voltage for all other components, within the same window. Peak latency was defined as the time point at which the peak amplitude occurred. For the 50% area latency, we first computed the area between the ERP difference wave and the zero-voltage line within the measurement window—using the area above zero for the P3 and below zero for the N170, MMN, N400, and ERN. The 50% area latency was then defined as the time point that divided this area into two equal halves. We final compared the changes for the four scores across different sampling rates.

### 2.5 Quantifying noise

Various in sampling rate will impact the noise. To quantify noise for different sampling rates, we computed the SME from the difference waveform for each ERP component for four scoring methods: mean amplitude, peak amplitude, 50% area latency and peak latency as previous studies (Luck et al., 2021; Zhang, Garrett, & Luck, 2024b; Zhang, Garrett, Simmons, et al., 2024; Zhang & Luck, 2023). For example, in the MMN experiment, we first computed a standard-minus-deviant difference wave, measured the amplitudes and latencies from the difference wave, and obtained the data quality for these scores separately. Note that the analyses were performed for a given ERP component at the maximal channel for that ERP as shown in Tab. 1.

SME values were computed from each individual participant for each score. For all the four scores, we used bootstrapping algorithm with replacement to estimate the SME from the corresponding difference wave directly (bootstrapping SME or bSME). The detailed process for bootstrapping is explained in Luck et al., 2021 and the simple MATLAB scripts are available at https://doi.org/10.18115/D58G91. In this study, we used 10000 iterations for each bSME value.

To obtain an overall measure of data quality across participants, we calculated the root mean square (RMS) of the single-participant SME values, referred to as RMS(SME). This was done by squaring each participant’s SME value, averaging these squared values across participants, and then taking the square root of the result. We used the RMS across participants rather than the mean because RMS is more directly related to effect size and statistical power (see the specific ratio for this in Luck et al., 2021). To estimate the standard error of the RMS(SME) values, we applied a bootstrapping procedure with 10,000 iterations.

### 2.6 SVM-based decoding analysis

For each ERP CORE paradigm, we decoded which of stimulus categories was given. For example, in the N400 experiment, we decoded whether the presented stimuli were related or unrelated. Each paradigm in the ERP CORE datasets contained two types of stimuli. Therefore, for each experiment, binary decoding was performed on the data of each participant at each time point. As demonstrated by previous studies (Trammel et al., 2023; Zhang & Luck, 2025), SVM outperformed other commonly used machine learning algorithms, such as linear discriminant analysis (LDA), we therefore employed SVM in the present study. Due to all paradigms in ERP CORE datasets being binary classification, we used the MATLAB function “fitcsvm()” to train the SVM classifier, and function “predict()” to test the decoder. To balance decoding accuracy with generalization, we employed the default regularization parameter (BoxConstraint =1) as previous study (Song et al., 2025; Zhang et al., 2025).

Referring to previous research reported (Carrasco et al., 2024; Song et al., 2025; Zhang et al., 2024), in this study, we used the leave-one-out 3-fold cross-validation to avoid overfitting. In each decoding analysis, for a given participant’s matrix, we divided it into separate matrices for each category based on *M* stimulus categories. To maximize decoding accuracy, we conducted decoding on the averaged ERPs, where each fold was associated with an average ERP waveform. Specifically, based on different random subsets of trials, generated 3 average ERPs for each category. For example, in the case of the N170 component, a total of 80 trials were collected for each of the Car and Face conditions prior to artifact rejection. To create more stable estimates, three averaged ERP sets were generated for each condition by randomly dividing the 80 trials into three subsets of approximately 26 trials each.

To perform decoding, the classifier was trained and tested using the three averaged ERP sets. In each iteration, two of the three averages were used for training, and the remaining one was used for testing. This three-fold cross-validation was repeated such that each ERP average served as the test set once at least. To enhance the reliability of the results, this entire procedure was repeated 100 times with different random assignments of trials to the averaged ERP sets in each iteration.

Decoding accuracy was defined as the proportion of correctly classified test cases. The chance level was set to the inverse of the number of stimulus categories, which was 1/2 (i.e., 0.5) for the ERP CORE datasets.

### 2.7 Effect size calculation

We assessed statistical power for both univariate and decoding analyses, using Cohen’s *d_z_* as the effect size metric.

For univariate analysis, Cohen’s *d_z_* represented the magnitude of the variance in the average of a single set of data. The method for calculating the Cohen’s *d_z_* of the four scores was to calculate the average value of each scoring method among different participants and then divide it by the standard deviation of each scoring method among different subjects. Then we estimated its standard error of mean using bootstrapping with 10,000 iterations.

Decoding accuracy varied over time, so to obtain a stable and interpretable estimate, we averaged the decoding accuracy within specific time windows defined by the standard univariate analysis ranges for each ERP component, as listed in Table 1. To assess statistical power, we calculated effect sizes using Cohen’s *d_z_*, which incorporates both the mean decoding accuracy and variability across participants. Specifically, Cohen’s *d_z_* was computed by subtracting the chance-level accuracy from the mean decoding accuracy across participants and dividing the result by the standard deviation across participants. The standard error of the effect size was estimated via bootstrapping with 10,000 iterations.

## 3 Results

In this section, we describe how sampling rate influences EEG data quality (quantified by RMS(SME)), conventional univariate ERP measures, and the decoding accuracy.

### 3.1 RMS(SME)

To evaluate how sampling rate influences EEG data quality, we computed RMS(SME) across a range of ERP components and dependent scores (mean amplitude, peak amplitude, 50% area latency, and peak latency). Lower values of RMS(SME) indicate better data quality as suggested by Luck (2021).

As shown in Fig. 2, across most ERP components and scores, sampling rate had minimal effect on RMS(SME) visually, particularly for mean amplitude and peak amplitude scores. ERP components such as P3, MMN, ERN, and N2pc exhibited highly similar RMS(SME) values across sampling rate ranging from 128 to 1024 Hz. This suggests that for amplitude-based metrics, data quality remains relatively stable event at lower sampling rates.

**Figure 2.**
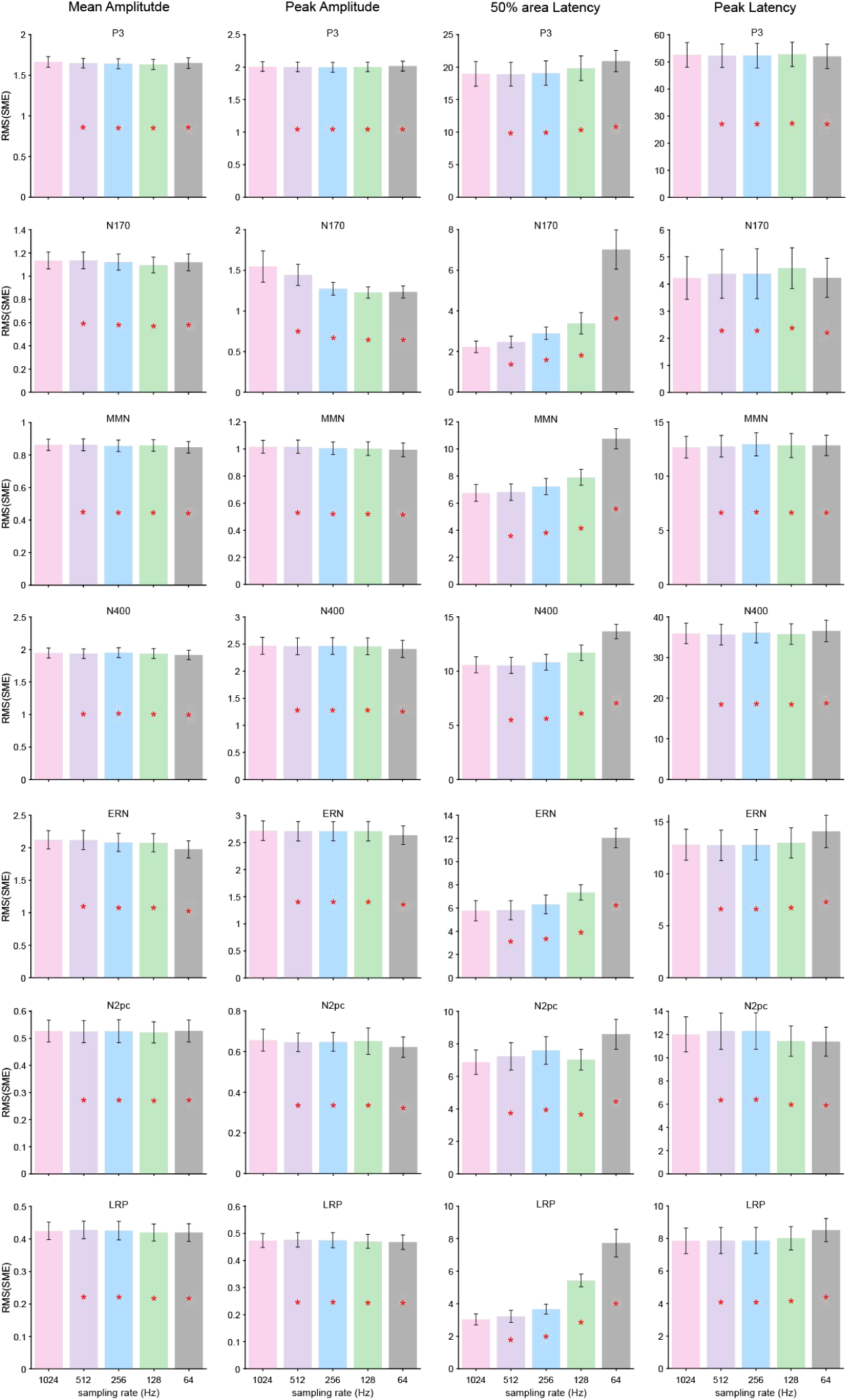
Impact of sampling rate on EEG data quality measured by root mean square of Standardized Measurement Error (RMS(SME)). The figure shows RMS(SME) for four commonly used ERP metrics—mean amplitude, peak amplitude, 50% area latency, and peak latency—across seven ERP components (P3, N170, MMN, N2, ERN, N2pc, LRP). Lower RMS(SME) values indicate better data quality (i.e., more reliable measurements). Each bar represents the standard error of mean (SEM) obtained by bootstrapping with 10,000 iterations. Asterisks mark cases in which SME differed significantly between a given sampling rate and the baseline one of 1024 Hz (according to a paired *t*-test). All p-values were corrected for multiple comparisons using the False Discovery Rate (FDR) across seven ERP components and all sampling rates.

However, latency-based measures, particularly 50% area latency, showed greater sensitivity to sampling rate (see Fig. 2). The N170, ERN, and LRP components demonstrated an increasing trend in RMS(SME) at lower sampling rate, indicating a decline in temporal precision. This was most pronounced for the 50% area latency of the N170, where RMS(SME) values increased sharply at 64Hz, suggesting a loss of reliability in latency estimates when temporal resolution is coarse.

We further compared the SME values obtained at 1024 Hz with those from other sampling rates using paired t-test. The analysis revealed a significant difference between the baseline condition at 1024 Hz and the other sampling rates for all scoring methods for all ERP components.

### 3.2 Conventional univariate analysis

We next examined how sampling rate influenced conventional ERP metrics (mean amplitude, peak amplitude, 50% area latency, and peak latency) across seven ERP components. The baseline sampling rate was set at 1024 Hz, and other rates ranged from 64 Hz to 512 Hz. Paired t-tests were used to assess statistical differences, indicated by asterisks for significance and “NS” for non-significant differences (Fig. 3). Additionally, we assessed how different sampling rates affect effect size (Cohen’s *d_z_*) across seven components for four commonly used score metrics (see Fig. 4).

**Figure 3.**
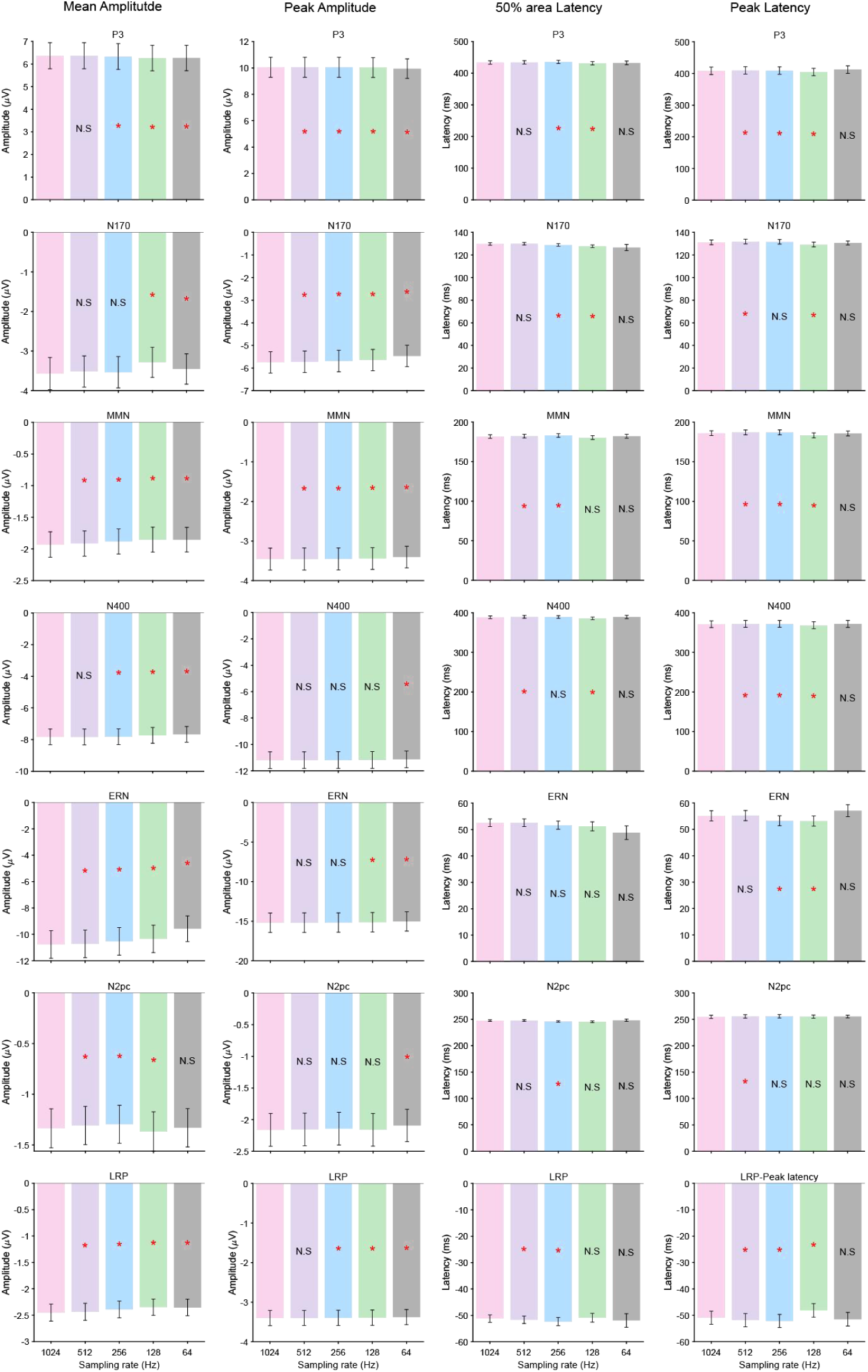
Effects of sampling rate on four ERP measurement scores across seven ERP components. The four columns represent (from left to right) mean amplitude, peak amplitude, 50% area latency, and peak latency. Each row corresponds to a different ERP component. Bars represent the average score at each sampling rate (64,128,256,512,1024Hz), with error bars indicating ±1 SEM. Asterisks mark cases in which scores differed significantly between a given sampling rate and the baseline one of 1024 Hz (according to a paired t-test). All p-values were corrected for multiple comparisons using the False Discovery Rate (FDR) across seven ERP components and all sampling rates.

**Figure 4.**
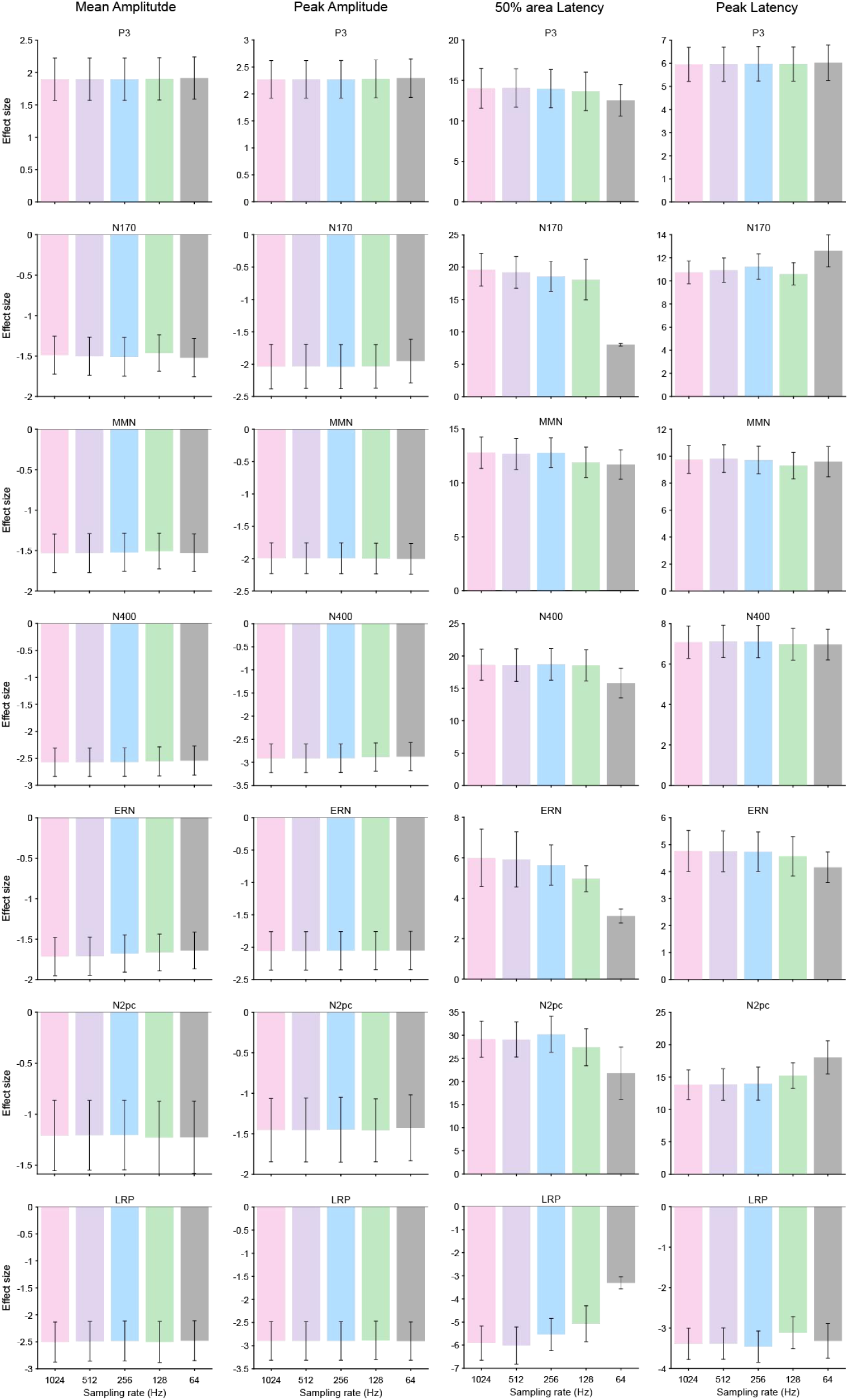
Effects of sampling rate on effect size of four ERP measurement scores across seven ERP components. The four columns represent the effect size for mean amplitude, peak amplitude, 50% area latency, and peak latency (from left to right) separately. Each row corresponds to a different ERP component. Bars represent the average score at each sampling rate (64, 128, 256, 512, 1024Hz), with error bars indicating ±1 SEM. The standard error of the effect size was estimated via bootstrapping with 10,000 iterations.

Fig. 3 presents the influence of sampling rate on different score metrics across seven ERP components. For most components, both mean and peak amplitudes showed some significantly variation with changes in sampling rate for most cases. However, peak amplitude for certain components (e.g., N400 and N2pc) remained largely stable across different sampling rates, indicating relative robustness of these measures to temporal resolution. Both peak latency and 50% area latency were largely preserved across sampling rates for most cases, suggesting that temporal resolution did not substantially affect the timing of ERP components.

Fig. 4 shows the effect size for four score metrics with different sampling rates across seven ERP components. Across all components and scoring methods, the effect sizes remained relatively stable at sampling rates of 128 Hz and above. However, a consistent and marked reduction in effect size was observed at the lowest sampling rate (64Hz), particularly for latency-based metrics (i.e., 50% area latency). This suggests that temporal precision at 128 Hz or lower may be insufficient to capture reliable latency differences, thereby attenuating sensitivity to experimental effects.

### 3.3 Binary Decoding

We further examined the impact of sampling rate on binary decoding accuracy and effect size across seven ERP components. Decoding was performed separately for each component, and average classification accuracy was computed across participants. Paired t-tests compared performance at 1024 Hz to other sampling rates.

As shown in Fig. 5, decoding accuracy was generally robust across sampling rates ranging from 64 Hz to 1024 Hz. For all ERP components, decoding performance remained virtually identical across sampling rates, as indicated by overlapping error bars and the absence of any systematic trend related to temporal resolution. Importantly, statistical analyses revealed no significant differences between the 1024 Hz baseline and any of the other sampling rates. These findings suggest that decoding performance for these components is largely insensitive to reductions in sampling rate.

**Figure 5.**
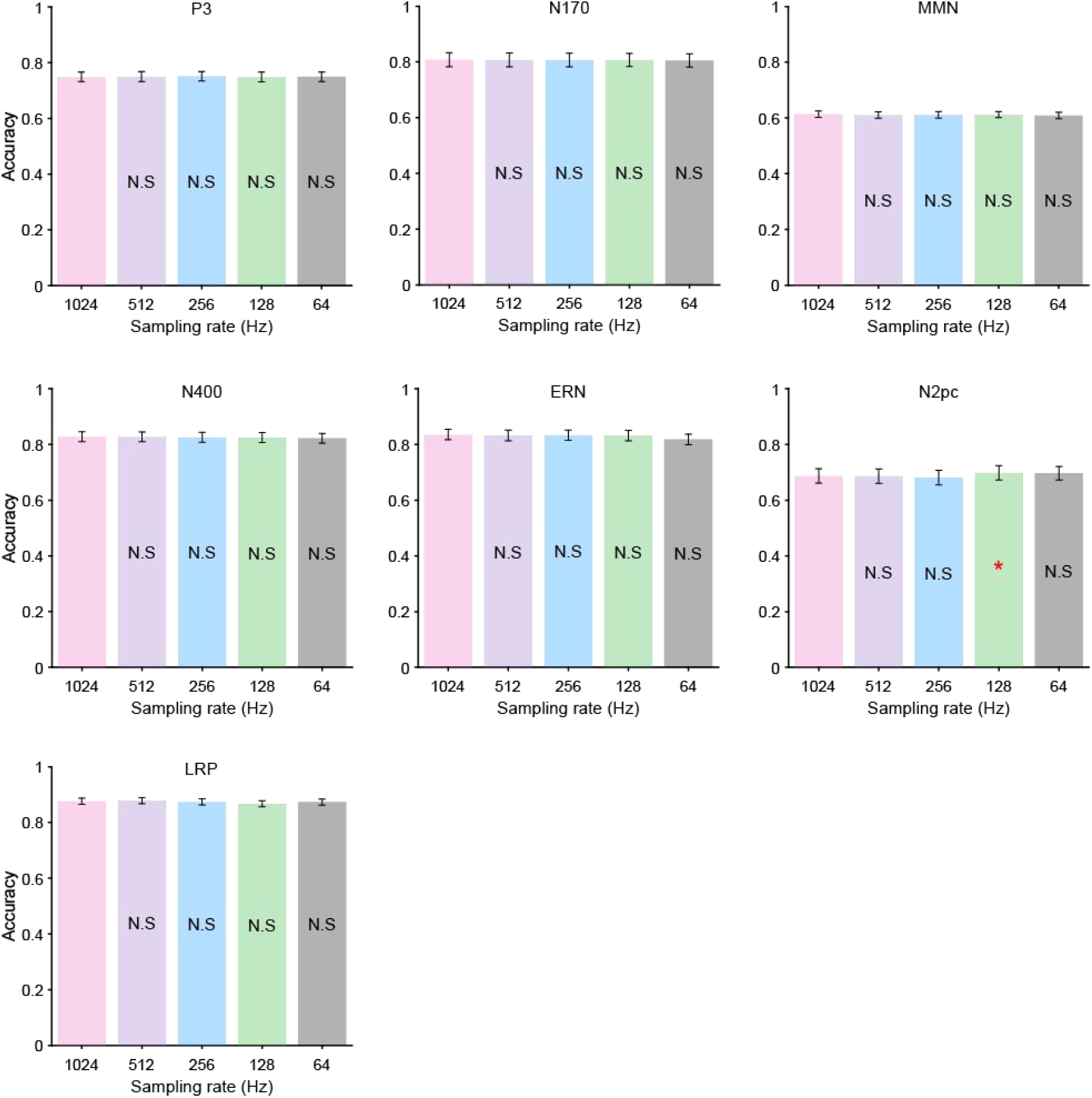
Effects of sampling rate on decoding accuracy across seven ERP components. Each row corresponds to a different ERP component. Bars represent the average score at each sampling rate (64,128,256,512,1024Hz), with error bars indicating ±1 SEM. Asterisks mark cases in which decoding accuracy differed significantly between a given sampling rate and the baseline one of 1024 Hz (according to a paired t-test). All p-values were corrected for multiple comparisons using the False Discovery Rate (FDR) across seven ERP components and all sampling rates.

Fig. 6 shows the effect size of decoding performance across varied sampling rates for seven ERP components. The results indicated that effect sizes were generally stable across sampling rates, with only small variations observed for few ERP components. For components such as P3, MMN, and LRP, effect sizes remained high and consistent across all sampling rates from 64 Hz to 1024 Hz, indicating robust decoding performance that was not strongly influenced by temporal resolution.

**Figure 6.**
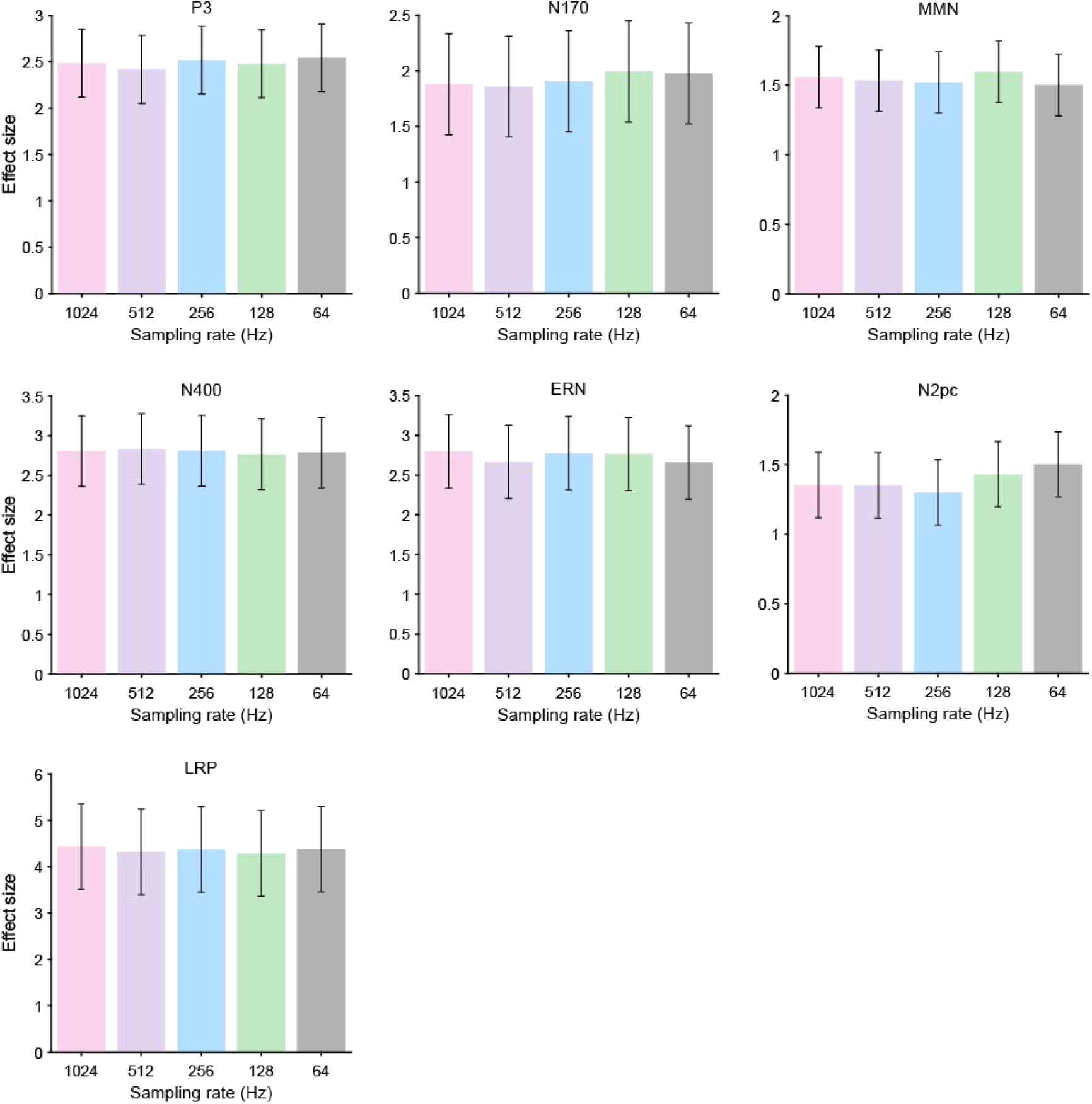
Effects of sampling rate on effect size of decoding accuracy across seven ERP components. Each row corresponds to a different ERP component. Bars represent the average score at each sampling rate (64,128,256,512,1024Hz), with error bars indicating ±1 SEM. The standard error of the effect size was estimated via bootstrapping with 10,000 iterations.

## 4. Discussion

In this study, we examined whether the downsampling has a significantly effect on the data quality, conventional ERP measurements, and decoding analysis. To our knowledge, it is first time to use a broad set of ERP datasets to assess the impact of sampling rate settings on these metrics. We found that both amplitude-based and latency-based data quality were significantly impacted by sampling rate. The 50% area latency measure exhibited lower data quality at reduced sampling rates compared with the other three measures. Traditional univariate analyses revealed that amplitude-based measures and their corresponding effect sizes were more sensitive to changes in sampling rate, whereas latency-based measures were comparatively less affected by variations in sampling rate. Furthermore, multivariate decoding presented strong stability, with both decoding accuracy and effect sizes showing minimal variation across sampling rates, even at the lower limit of 64 Hz.

A modest sampling rate might have been expected to improve data quality by reducing trial-to-trial variability; however, it did not significantly reduce the SME for amplitude-based metrics across most of the seven ERP components examined in this study. An exception was found for the peak amplitude of the N170, where a decrease in sampling rate was associated with reduced SME, indicating an improvement in data quality. In contrast, latency-based metrics were more sensitive to changes in sampling rate, particularly the 50% area latency. It is possible that downsampling may significantly impact on data quality in experimental paradigms with substantially lower signal-to-noise ratios than those used in the current datasets, which was derived from highly cooperative college student participants.

For conventional univariate analyses, the impact of sampling rate varied across ERP components and scoring methods. Significant differences were observed between the 1024 Hz baseline and other sampling rates for most amplitude estimates, whereas latency-based measures were generally less affected by changes in sampling rate. These findings are broadly consistent with previous studies indicating that moderate downsampling does not substantially reduce the statistical power of amplitude-based measures for some ERP components (e.g., peak amplitude for N400; Clayson et al., 2013). Importantly, the present results extend this evidence by showing that latency-based indices of some components, such as the ERN, can also tolerate moderate downsampling. Together, these observations suggest that the impact of sampling rate on conventional univariate analyses depends on both the ERP component under consideration and the scoring method employed.

Notably, multivariate decoding performance was highly robust across all tested sampling rates from 64 to 1024 Hz. Both decoding accuracy and corresponding effect sizes remained stable across ERP components, with no systematic trend of degradation at lower sampling rates. This finding suggests that decoding methods, which leverage spatiotemporal patterns rather than isolated peak values, are relatively robust to temporal downsampling. This robustness likely arises from the distributed nature of the information used by multivariate classifiers, which can tolerate coarser temporal resolution as long as spatial and overall temporal structure are preserved (Grootswagers et al., 2017). The resilience of decoding approaches even at 64 Hz suggests that under typical experimental designs, decoding analyses may not require high-frequency sampling to achieve meaningful classification performance. This likely explains the widespread use of lower sampling rates, such as 50 Hz, in ERP decoding studies (Bae & Chen, 2024; Carrasco et al., 2024; Zhang et al., 2024; Zhang & Luck, 2025). However, for tasks requiring precise timing (e.g., fast sequences or fine-grained temporal generalization), higher sampling rates may still be advisable (A.-S. Li et al., 2024; Y. Li et al., 2022; Tautvydaitė & Burra, 2024).

Our findings have practical implications for ERP research and related signal processing decisions. For researchers focusing on amplitude- or latency-based ERP measures, careful consideration of sampling rate is warranted. In contrast, when applying multivariate decoding techniques, downsampling to 128 Hz or even 64 Hz can offer substantial reductions in data storage and computational demands with minimal impact on performance. Given the growing use of portable and wireless EEG systems with limited sampling rates (He et al., 2023), our findings provide empirical support for their effectiveness in capturing reliable ERP and decoding measures, provided that the sampling frequency remains above the minimum thresholds of 512 Hz for ERP and 64 Hz for decoding analyses.

This study focused on a representative set of ERP components recorded under typical experimental conditions from neurotypical adults. Generalization to other paradigms, higher-noise settings, infants, or clinical populations should be approached cautiously (Ashton et al., 2022; Marsicano et al., 2024; Ng et al., 2022). Future work could extend these findings by systematically evaluating how sampling rate interacts with other preprocessing steps (e.g., filtering, interpolation), stimulus timing accuracy, or decoding architectures. Additionally, time-resolved decoding approaches (e.g., temporal generalization matrices) might be more sensitive to downsampling than the component-wise analyses employed here (King & Dehaene, 2014). Furthermore, the present findings may not readily extend to more complex decoding approaches based on deep learning architectures, such as convolutional neural networks, recurrent neural networks, or deep belief networks (Al-Saegh et al., 2021; Craik et al., 2019).

### CRediT authorship contribution statement

Guanghui Zhang:Writing – review & editing, Writing – original draft, Visualization, Validation, Supervision, Software, Resources, Methodology, Conceptualization; Xinran Wang:Writing – review & editing, Writing – original draft, Visualization, Validation, Software, Resources, Methodology, Investigation; Ying Xin: Writing – review & editing, Writing – original draft, Visualization, Validation; Fengyu Cong: Writing – review & editing, Writing – original draft, Visualization, Validation; Weiqi He: Writing – review & editing, Writing – original draft, Visualization, Validation, Supervision; Wenbo Luo: Writing – review & editing, Writing – original draft, Visualization, Validation, Supervision.

## Declaration of competing interest

None.

## Data availability

The data and analysis scripts are available at: https://osf.io/v5suh/.

## Footnotes

1 This identical pre-filtering across sampling-rate conditions prevents frequencies above each condition’s Nyquist limit from being aliased into the passband during down-sampling. Because the ERP components analyzed here are distributed well below 20 Hz, the 20 Hz cutoff preserves the signals of interest while eliminating potential spectral leakage; consequently, comparisons across sampling rates reflect sampling-rate effects rather than differences in aliased high-frequency content.

## Notes

**Funding Information:** This study was funded by the National Natural Science Foundation of China (No. 32471107), the Scientific Research and Innovation Team of Liaoning Normal University (No.25GDL004), and the Natural Science Foundation of Liaoning Province (No. 2025-BS-0780).

### Competing Interest Statement

The authors have declared no competing interest.

